# Lyn controls chemotaxis and motility of CLL cells via phosphorylation of ROR1

**DOI:** 10.1101/2020.05.29.124156

**Authors:** Zankruti Dave, Olga Vondálová Blanářová, Štěpán Čada, Pavlína Janovská, Nikodém Zezula, Martin Běhal, Kateřina Hanáková, Sri Ranjani Ganji, Pavel Krejci, Kristína Gömöryová, Helena Peschelová, Michael Šmída, Zbyněk Zdráhal, Šárka Pavlová, Jana Kotašková, Šárka Pospíšilová, Vítězslav Bryja

**Author notes:** **Corresponding author** (*): Prof. Vítězslav Bryja, PhD. Department of Experimental Biology, Faculty of Science, Masaryk University, Kotlářská 2, 611 37, Brno Czech Republic.

## Abstract

Chronic lymphocytic leukemia (CLL) and mantle cell lymphoma (MCL) are malignancies characterized by the dependence on B-cell receptor (BCR) signaling and by the high expression of the cell surface receptor ROR1. Both, BCR and ROR1 are therapeutic targets in these diseases and the understanding of their mutual cross talk is thus of direct therapeutic relevance. In this study we analyzed the role of Lyn, a kinase from the Src family, as a mediator of the BCR-ROR1 crosstalk. We confirm the functional interaction between Lyn and ROR1 and demonstrate that Lyn kinase efficiently phosphorylates ROR1 in its kinase domain and aids the recruitment of an E3 ligase c-CBL. The absence of Lyn in Lyn KO Maver-1 cells produced by CRISPR-Cas9 resulted in the increased ROR1 cell surface levels and deregulated migratory properties. Similar correlations between ROR1 surface dynamics, levels of active Lyn and chemotactic properties were confirmed in primary CLL samples. Our data establish Lyn-mediated phosphorylation of ROR1 as a point of crosstalk between BCR and ROR1 signaling pathways.

## Introduction

The ROR protein family comprises of ROR1 and ROR2, which are both type 1 transmembrane receptors. Upon discovery, ROR proteins were referred to as orphan receptors on account of the lack of identity of their ligands. However, subsequent studies identified their ligands to be the Wnt proteins, mostly Wnt-5a protein^1,2^. Wnt-5a/ROR pathway is an essential signaling pathway that controls cell polarity and migration during embryonic development and tissue homeostasis^3,4^ During embryonic development, RORs are highly and uniformly expressed, most prominently in the skeletal and neural tissues, but postnatally their expression becomes highly restricted^5^.

Interestingly, ROR1 or ROR2 upregulation has been observed in many cancers: ROR1 is upregulated in solid tumors or hematologic malignancies while ROR2 is overexpressed in osteosarcomas or renal cell carcinomas^6^. High expression of ROR1 is typical for some B-cell lymphomas such as mantle cell lymphoma (MCL)^7^ and chronic lymphocytic leukemia(CLL)^8,9^. CLL is a form of hematologic cancer which is manifested as a steady accumulation of mature CD5^+^ B-cells in the bone marrow, lymphoid tissues and peripheral blood. It is the most common form of adult leukemia in the western hemisphere, with an incidence of 5 per 100 000 each year and an average median age of on-set around 70 years. Most of the CLL cases remain asymptomatic for a long time, in which case therapeutic intervention is not necessary. However, part of the CLL cases progress rapidly, require treatment and their overall life expectancy is decreased^10^.

CLL cells are in most cases highly ROR1 positive^1,8^ and there are currently several therapies in development that target ROR1^11,12^. Another typical feature of CLL is the dependency on the B-cell receptor signaling (BCR) pathway that promotes survival and proliferation of CLL cells^13,14^. Modern treatments in CLL are thus designed to target the BCR pathway components of which some examples include: Bruton tyrosine kinase (BTK) inhibitor ibrutinib, and PI3K targeted by idelalisib^10^. Importantly, there are several pieces of evidence that propose a mutual crosstalk between ROR1 and BCR signaling^15–17^. Given the importance of both pathways in the novel therapeutic strategies for CLL/MCL, identification of the molecular basis of such crosstalk would be of utmost importance.

In this study we have analyzed the role of Lyn, a kinase from the Src family, as a candidate for such a function. It is the predominant Src family kinase in lymphoid cells and it plays a dual role as the positive as well as negative regulator of the BCR pathway^18^. We focused on Lyn because of prior studies which identified the important role of Src in the regulation of ROR1 or ROR2 in other experimental systems^19,20^. Further, Lyn as an important component of BCR pathway, has been found to be over expressed in CLL patients^21^. We were able to confirm the mechanistic and functional interaction between Lyn and ROR1 in several cell types including MCL cell line Maver-1 and primary CLL. Our study establishes Lyn-mediated phosphorylation of ROR1 as another point of crosstalk between BCR and ROR1 signaling pathways.

## Materials and Methods

### Cell culture

All cell lines used in the experiments were grown at 37°C and 5% CO_2_. HEK-293T wild type cells (ATCC-CRL-11268, LGC Standards, Manassas, VA) were cultured in DMEM medium (Thermo Fischer Scientific, USA) supplemented with 10% FBS (#10270, Gibco, Thermo Fisher Scientific) and 1% Penicillin/Streptomycin (#sv30010, HyClone, GE Healthcare, Chicago, IL). The mantle cell lymphoma cell line Maver-1 (#ACC 717, DSMZ Gmbh, Braunschweig, Germany) was cultured in HyClone™ RPMI 1640 medium (GE Healthcare) supplemented with 10% heat-inactivated FBS and 1% Penicillin/Streptomycin.

### Primary CLL samples

Patient samples were obtained from Dept. of Internal Medicine – Hematology and Oncology, University Hospital Brno. B cells were isolated from the peripheral blood of CLL patients undergoing monitoring and treatment at the hospital, as described here^22^. CLL samples were obtained after written informed consent in accordance to the Declaration of Helsinki and by following protocols approved by the ethical committee of the University Hospital, Brno. Primary CLL cells were grown in HyClone™ RPMI 1640 medium (GE Healthcare) supplemented with 10% heat-inactivated FBS and 1% Penicillin/Streptomycin at 37°C and 5% CO_2_.

### Transfections, treatments and plasmids

Transient transfections of HEK-293T cells were carried out using polyethyleneimine (PEI 1 mg/ ml); for transfections in a 10 cm plate, a total of 6 μg of DNA and for the 24 well plate a total of 0.2 μg of DNA per well was used. The DNA:PEI ratios were kept at 1:3 (w/v) in all cases. All the ROR1 plasmids, apart from the *ROR1-v5-his*, were provided by Prof. Paolo Comoglio and have been described previously^20^. All the Lyn expression plasmids were a kind gift from Dr. Naoto Yamaguchi^23^. Dasatinib (#sc-218081, Santa Cruz Biotechnology, Santa Cruz, CA) was used to inhibit the kinase activity of Lyn. Further details are provided in the online Supplementary Materials and Methods.

### Immunoprecipitation, western blotting and immunocytochemistry

An extensive description is provided in the online Supplementary Materials and Methods but, in brief, for the immunoprecipitation experiments, transfected cells were first washed in cold PBS and then lysed in cold 0.5% NP-40 Lysis buffer and kept at 4°C. Prior to lysis, the buffer was supplemented with 1mM Na_3_VO_4_, 1mM DTT, 1mM NaF and with cOmplete™ protease inhibitor cocktail and phosphatase inhibitor cocktail set II (Merck, Kenilworth, NJ). For the western blotting, samples were loaded on 8% gels and separated by SDS-PAGE followed by the transfer done on to Immobilon-P^®^ (Merck) PVDF membranes at 106 V for 75 minutes.

For immunocytochemistry, HEK-293T cells were grown on glass coverslips, transfected, and immunostained. The images were taken using Leica SP8 confocal microscope.

### Generation Lyn KO Maver-1 cell line

The knockout of Lyn gen in Maver-1 was performed using Crispr/Cas9 system. Selection of the Lyn KO was based on the results of western blot and confirmed using next generation sequencing of PCR product as described here^24^. A detailed description can be found in the online supplementary methods.

### Quantitative real-time PCR

Quantitative real-time qPCR was used to assess the relative gene expression of ROR1 in WT and Lyn KO Maver-1 cells. Data were analyzed by Delta-Delta C_T_ method and are shown as 2^-ΔΔCT^^25^. A detailed description can be found in the online supplementary methods.

### Transwell migration assay

Cell migration assays were carried out using HTS Transwell 24-well plates with a 5 μm pore size polycarbonate membranes (Corning, New York, NY). 0.2 x 10^6^ WT or Lyn KO Maver-1 cells or 1 x 10^6^ primary CLL cells were seeded into the transwell upper inserts while media were supplemented with chemokine CCL19 (#361-MI-025, R&D Systems, Minneapolis, MN) at 200 ng/mL or 0.1% BSA in PBS (control) in the lower chamber. After 3 h (Maver-1 cells) or 6 h (primary CLL cells) at 37°C with 5% CO_2_, the cells in the lower chamber were collected and counted using the Accuri C6 Flow Cytometer (Becton Dickinson, Franklin Lakes, NJ).

### BCR Stimulation assay

Maver-1 cells (1x 10^6^) were subjected to BCR stimulation for 4 min at 37°C, with 10 μg/mI F(ab’)_2_ Anti-human anti-IgM-UNLB (#2022-01, Southern Biotech, Birmingham, AL) resuspended in PBS. The control samples were treated with 0.1% BSA in PBS. Post stimulation the cells were immediately spun at 200 g for 5’ and washed with PBS. The cells were finally pelleted and resuspended in 1x Laemmli buffer. The samples were sonicated for 15 s and further subjected to Western blotting.

### Flow cytometric analysis of surface expression of CCR7 and ROR1

Cells either from culture or from transwell assay were washed in PBS and incubated in 2% FBS in PBS with anti-CCR7-FITC (1:25, #561271, BD Biosciences) and anti-ROR1-APC (1:25, #130-119-860, Miltenyi Biotec, Bergish Gladbach, Germany) antibodies on ice for 20 minutes. The cells were washed and resuspended in PBS and analyzed using Accuri C6 Flow Cytometer (BD Biosciences). Data were analyzed using NovoExpress (ACEA Biosciences, Inc., San Diego, CA) and presented as a median fluorescence intensity (MFI index) or as a fold of MFI of wt cells (Maver Lyn KO cells) or as a ratio of MFI of cells from lower and upper compartment.

### Mass spectrometry

Unbiased identification of ROR1 phosphorylation sites and ROR1 interaction partners was performed by mass spectrometry. A detailed description can be found in the online supplementary methods.

### Statistics

All statistical tests were performed using GraphPad Prism software 6.0 (GraphPad Prism Software, Inc., San Diego, CA). Number of replicates, format of data visualization and statistical tests used for comparison are indicated in the individual figure legends.

## Results

### Lyn interacts with the Wnt-5a receptor ROR1

ROR1 has been reported earlier to interact with the members of the Src kinase family^20,26^. In order to test if this holds true for ROR1 and Lyn, we overexpressed both proteins in HEK-293T cells and performed immunoprecipitation experiments. We observed a strong and specific pulldown of ROR1 by anti-Lyn antibody and vice versa (**Fig. 1a** and **1b**). In order to visualize this interaction intracellularly we used immunocytochemistry. Overexpressed ROR1 and Lyn co-localized in the cell membrane and filopodia of HEK-293T cells (**Fig. 1c)**. Next, we attempted to identify the domains of ROR1 involved in its interaction with Lyn. For this purpose, we used a set of ROR1 deletion mutants^20^ (**Fig. 1d**). Lyn was able to interact with the WT ROR1 as well as with the ROR1 lacking the C-terminal tail formed by two Ser/Thr-rich and one Pro-rich (PRD) domains. Further deletion of the complete intracellular domain of ROR1 abolished the binding to Lyn (**Fig. 1e**). These results indicated that the ROR1 kinase domain, and/or nearly adjacent regions, represent a crucial interaction interface for Lyn.

**Figure 1:**
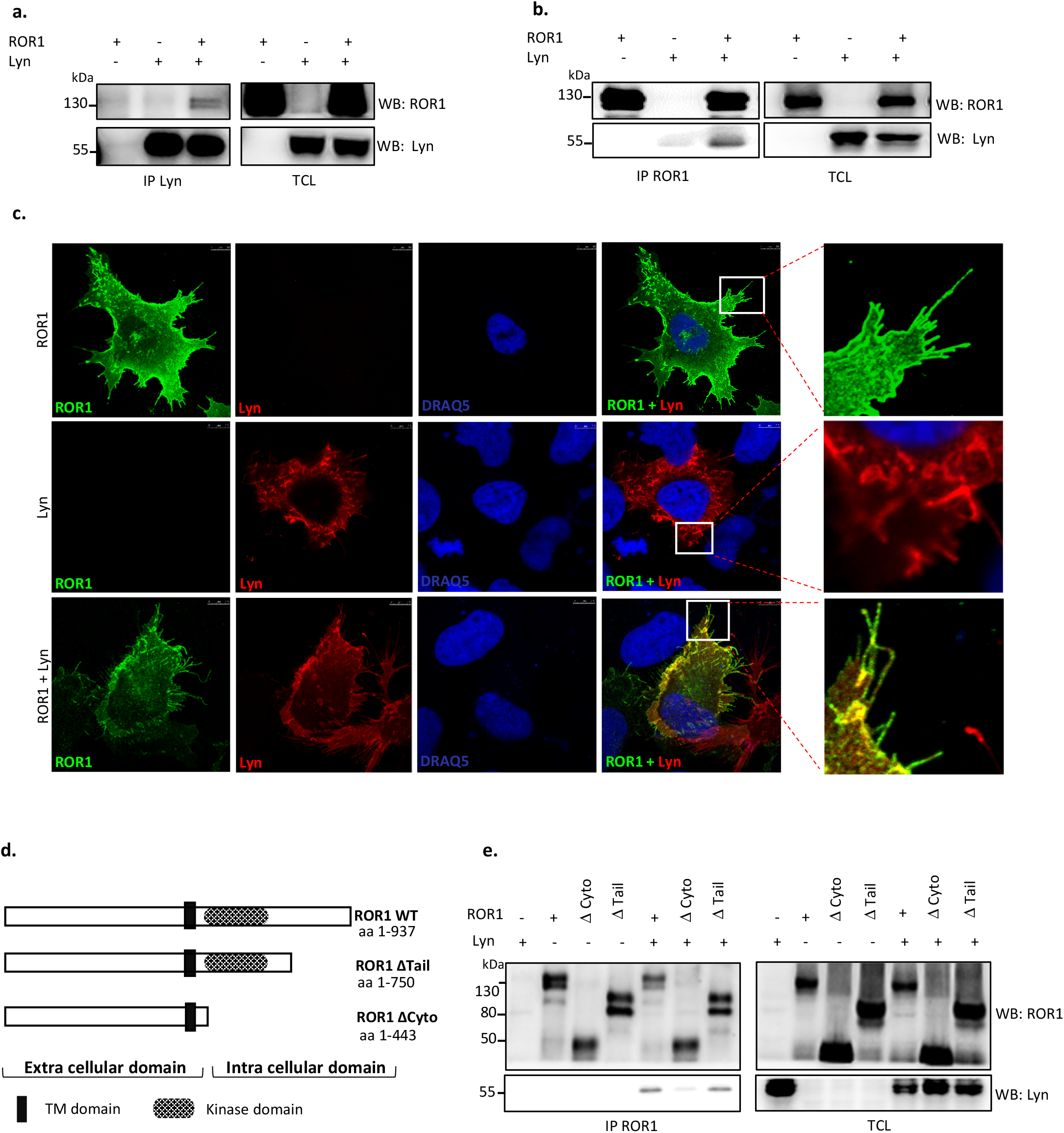
Lyn interacts with ROR1. a, b) Lyn and ROR1 were overexpressed in HEK-293T cells. co-IP and Western blot analysis showing a pull-down of ROR1 when the lysates were immunoprecipitated with Lyn (a) and a pull-down of Lyn when the lysates were immunoprecipitated with ROR1 (b). c) Representative images of immuno-cytochemistry analysis of HEK-293T cells overexpressing ROR1 (green) and Lyn (red) in the indicated combinations. Co-localization of ROR1 and Lyn is observed at the membrane. Scale bar 7.5 μm. d) Scheme of ROR1 mutants used for domain mapping. e) Lyn was co-expressed with the ROR1 intracellular deletion mutants in WT HEK-293T cells. Immunoprecipitation was done using ROR1 as the bait. WB – Western blotting, IP – immunoprecipitation, TCL – total cell lysate. Results in all panels are representative of at least 3 biological replicates.

### Lyn phosphorylates ROR1

Lyn is a tyrosine kinase and as such we wanted to test whether ROR1 represents its substrate. ROR1, a member of the receptor tyrosine kinase family, has a considerable number of tyrosine (Y) residues, which can be phosphorylated. Thus, we utilized a set of Lyn plasmids [23] (schematized in **Fig. 2a)** to test this hypothesis. Lyn activity is regulated by phosphorylation: Phosphorylation of the Y508 at its C-terminus keeps Lyn inactive and dephosphorylation of this site is necessary for the activation of Lyn. On the other hand, (auto)phosphorylation at Y396 turns it into an active kinase^27,28^. We used the following Lyn variants: WT Lyn; kinase domain deleted (Δ, aa 1-298) mutant lacking a significant portion of the kinase domain; kinase active Lyn (KA, aa 1-506) lacking the C-terminal inhibitory Y508 and allowing the constant activation of Lyn; and kinase dead Lyn (KD, aa 1-506, K275A) Lys->Ala mutation in the ATP binding pocket rendering Lyn kinase dead.

**Figure 2:**
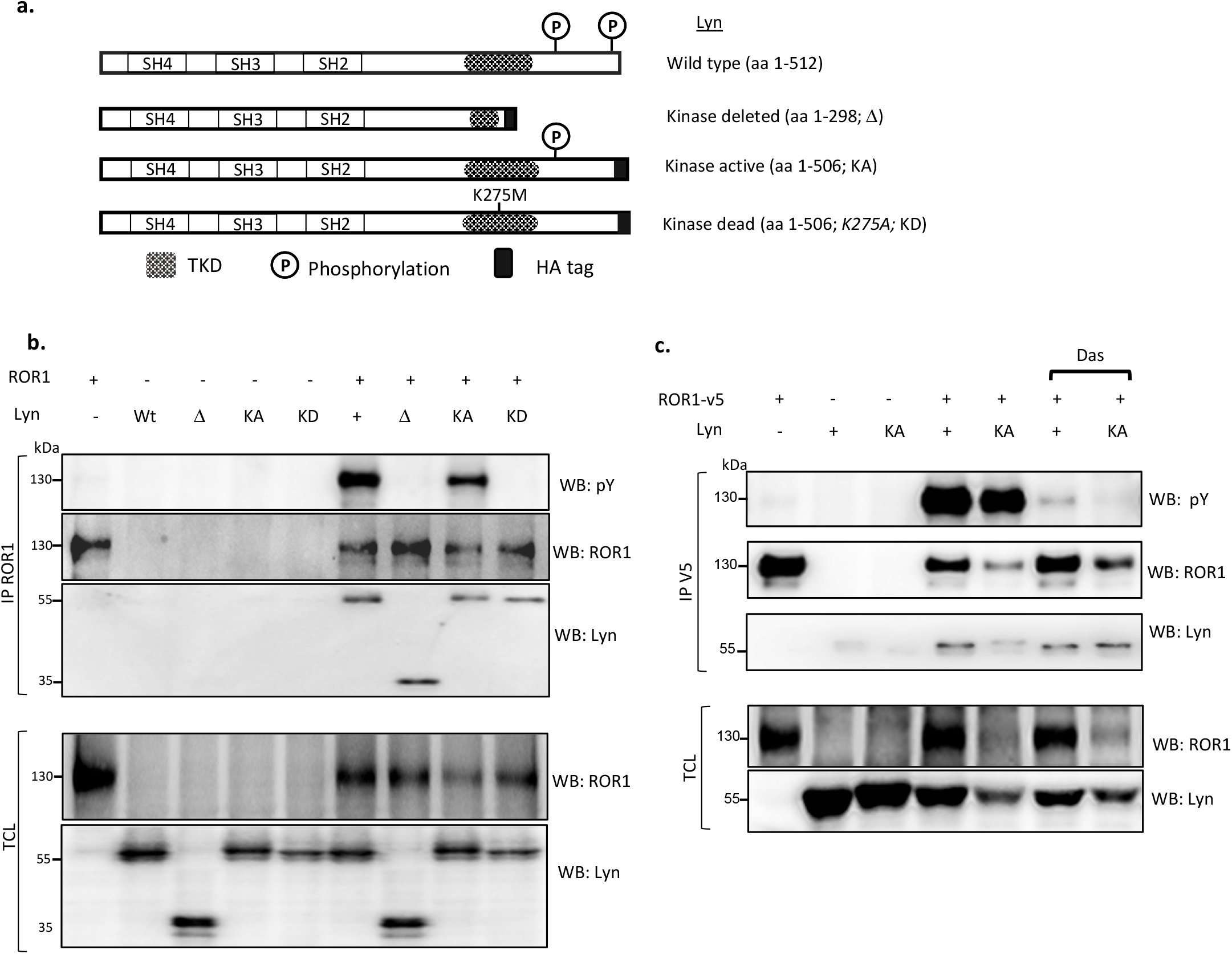
Lyn phosphorylates ROR1. Indicated combinations of Lyn and ROR1 plasmids were overexpressed in HEK-293T cells. ROR1 was immunoprecipitated and the binding of Lyn and Y phosphorylation was assessed by Western blotting. a) General scheme of the Lyn mutants used in b/c. b) Only the Lyn mutants with the intact kinase activity were able to phosphorylate ROR1 on its tyrosine residues. c) Small molecule inhibitor of Lyn, Dasatinib (Das, 0.2 μM), did not affect the interaction of ROR1 and Lyn, however it did block the ability of Lyn to phosphorylate ROR1. WB – Western blotting, IP – immunoprecipitation, TCL – total cell lysate. Results in all panels are representative of at least 3 biological replicates.

WT Lyn and kinase-active (KA) Lyn triggered a strong phosphorylation of ROR1 that could be detected by phospho-tyrosine (pY) specific antibody of immunoprecipitated ROR1 (**Fig 2b)**. In contrast, Lyn Δ and Lyn KD failed to phosphorylate ROR1 (**Fig. 2b**), which suggests that the phosphorylation depends on Lyn kinase activity. Interestingly, all 4 different forms of Lyn efficiently interacted with ROR1 (**see Fig. 2b**; IP ROR1, WB: Lyn). To further corroborate the analysis, we pharmacologically inhibited the kinase activity of Lyn by Dasatinib, a pan-Src family inhibitor^29^. Dasatinib did not interfere with the interaction between ROR1 and Lyn WT or Lyn KA, however, it did interfere in the phosphorylation of ROR1 as seen by the substantial decrease of tyrosine phosphorylation on ROR1 (**Fig. 2c)**.

### Identification and validation of ROR1 tyrosine residues phosphorylated by Lyn

In order to identify ROR1 residues that are phosphorylated by Lyn, we immunoprecipitated ROR1 in presence and absence of Lyn from HEK-293T cells and subjected them to mass-spectrometry analysis of phosphorylation(s). The experimental design is schematized in **Fig. 3a.** Proteomic analysis detected phosphorylated tyrosines only when Lyn was co-expressed. In total 6 tyrosine residues - Y459, Y645, Y646, Y666, Y828, and Y836 - were found phosphorylated exclusively in the presence of Lyn (**Fig. 3b).** Three residues are part of ROR1 tyrosine kinase domain (TKD) and two of those – Y645 and Y646 overlap with the residues reported to be phosphorylated by Src^20^. In order to decipher which of those residues are functionally important, we generated ROR1 individual point mutants as well as Y645/Y646 double mutant (schematized in **Fig. 3c).** Even though all tested mutants showed decreased levels of phosphorylation, the double Y645/Y646 was clearly the most deficient **(Fig 3d)**. This suggests that Y645 and Y646 are the most critical residues phosphorylated by Lyn.

**Figure 3:**
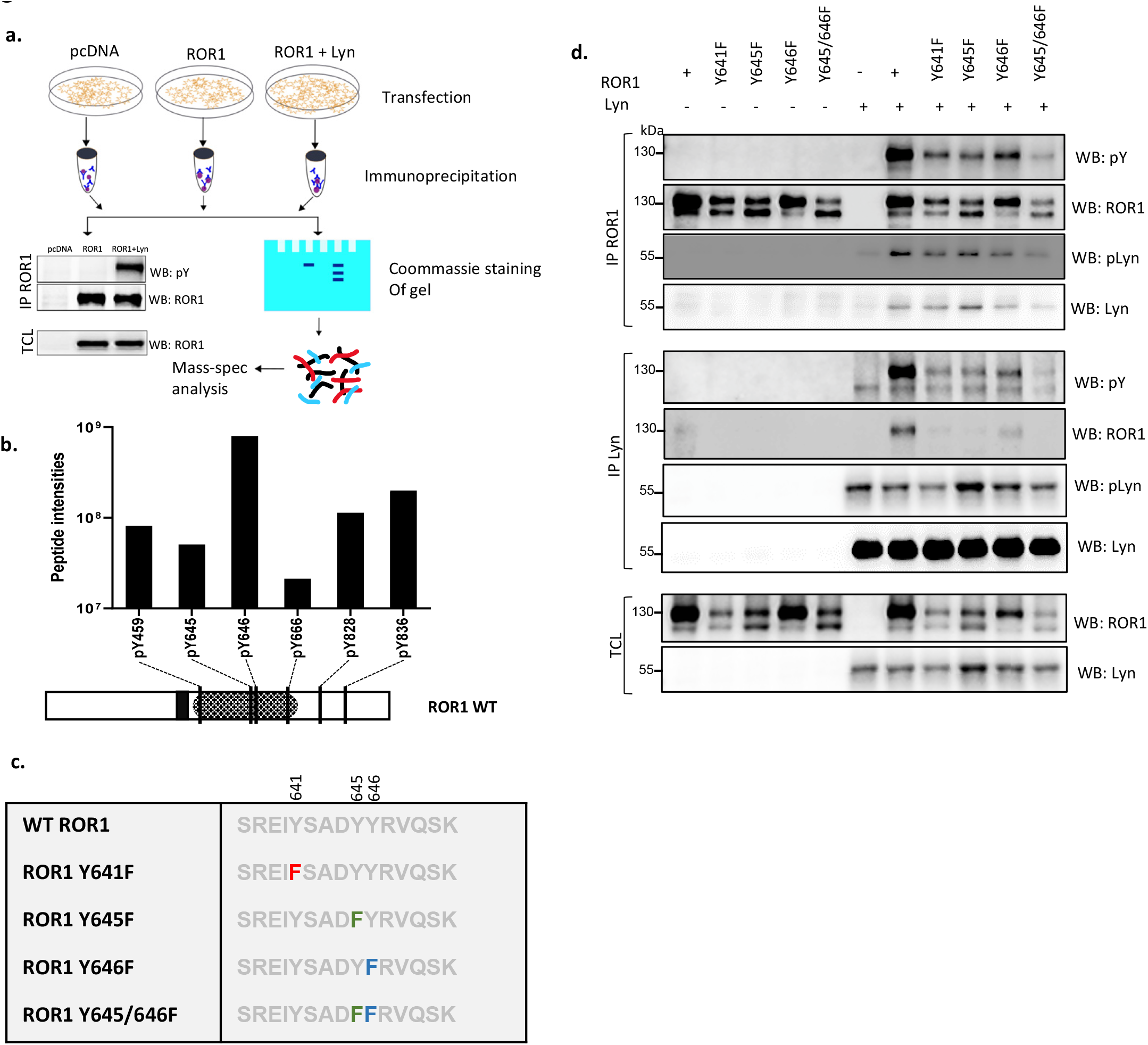
Mapping of the ROR1 residues phosphorylated by Lyn. **a)** Scheme of the experimental set-up for mass-spec analysis of ROR1 phosphorylation. Indicated combinations were transfected in HEK-293T cells. ROR1 was immunoprecipitated, separated on SDS-PAGE and bands corresponding to ROR1 were analyzed by MS/MS. **b**) ROR1 tyrosine (Y) residues that were found phosphorylated only when Lyn was coexpressed as identified by MS/MS analysis. The phospho-peptide signal intensity (up) and the position (bottom) of each detected phospho-tyrosine is presented. **c)** Schematics of the point mutants of ROR1 made for the validation experiments. **d**) Phosphorylation analysis of the point mutants showed that Y645/646F ROR1 mutation almost completely eliminated the phosphorylation by Lyn. WB – Western blotting, IP – immunoprecipitation, TCL – total cell lysate. Results in d are representative of at least 3 biological replicates.

### Lyn-induced phosphorylation of ROR1 induces recruitment of E3 ligase c-CBL

Phosphorylation at tyrosines is a well described signaling event with various functional consequences^30^. Often, phosphorylated tyrosines serve as molecular motifs recognized by downstream proteins containing SH2 domain. We thus hypothesized that ROR1 phosphorylation by Lyn will lead to the recruitment of further signal regulators. In order to address this question, we decided to identify ROR1 interaction partners induced by Lyn phosphorylation using unbiased immunoprecipitation coupled to mass spectrometry (IP/MS). We overexpressed ROR1 either alone or with WT or Lyn KD; pcDNA and Lyn WT-only transfected cells served as a control. Design of the experiment is schematized in **Fig. 4a**. IP/MS analysis identified 13 proteins that were uniquely detected as binding partners of ROR1 phosphorylated by Lyn (**Fig. 4b**). Among the hits (**Fig. 4c**), c-Casitas B lineage lymphoma (c-CBL) protein attracted our attention. c-CBL is an E3 ligase that recognizes pY motifs^31^ and often downregulates its targets by triggering them for degradation^32^ or endocytosis^33^. Of note, it is a known binding partner of Lyn^32^, as well as its substrate^34^.

**Figure 4:**
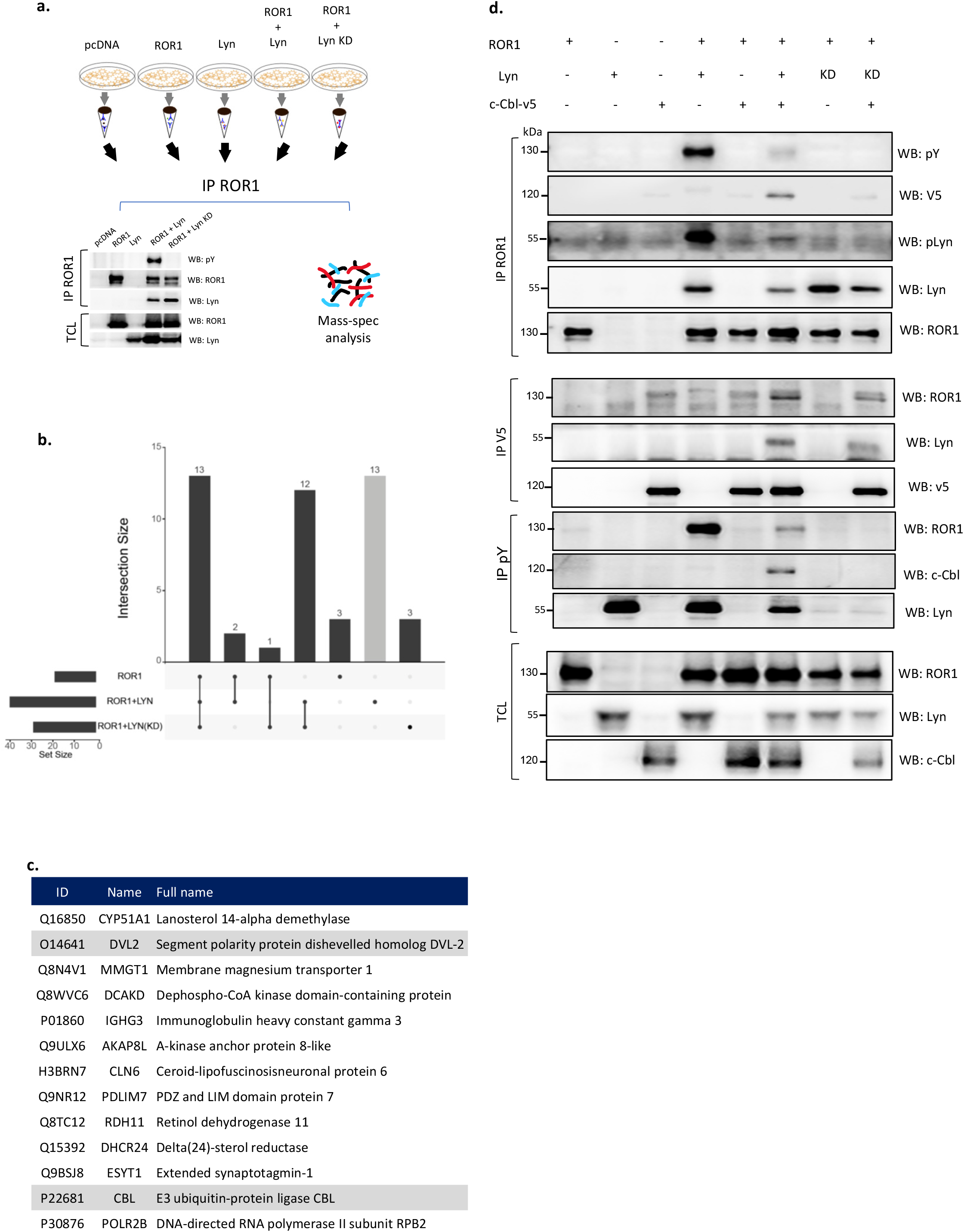
Lyn induced phosphorylation of ROR1 triggers interaction with the E3 ligase c-CBL. **a)** Scheme of the experimental set-up for the analysis of ROR1 interacting partners by MS/MS. ROR1 and Lyn WT and Lyn KO were overexpressed in HEK-293T. ROR1 was immunoprecipitated and the protein composition of the pulldown was analyzed by MS/MS. **b)** Upset plot demonstrating the numbers of proteins identified as ROR1 interactors in ROR1, ROR1+WT Lyn and ROR1+Lyn kinase dead (KD) conditions. Only proteins absent in the control pulldowns (pcDNA and Lyn expression) were considered. **c)** List of ROR1 interactors identified only when phosphorylated by Lyn. **d)** Analysis of the interactions and phosphorylation status of ROR1 and c-CBL. Indicated combinations were overexpressed in HEK-293T, immunoprecipitated (IP) as indicated and subsequently analysed by WB. Lyn promotes the interaction of ROR1 with c-Cbl. WB – Western blotting, IP – immunoprecipitation, TCL – total cell lysate. Results in d are representative of at least 3 biological replicates.

We overexpressed the combinations of c-CBL, ROR1 and Lyn plasmids in HEK-293T cells and performed a set of immunoprecipitation assays. In line with the mass spectrometry data, c-CBL efficiently interacted with ROR1 only when WT Lyn was present. The interaction between ROR1 and c-CBL was dependent on the Lyn-mediated phosphorylation of ROR1 since in the presence of the Lyn KD the binding to ROR1 was reduced **(Fig. 4d,** IP ROR1, WB V5, lanes 6 vs. 8). Lyn was a part of the complex since it was pulled down both by ROR1 and c-CBL **(Fig. 4d,** IP ROR1 & IP V5). Of note, co-expression of c-CBL clearly attenuated the phosphorylation of ROR1 by Lyn and level of active Lyn itself (**Fig. 4d**, IP ROR1 and IP pY, WB ROR1, Lyn and pY). Altogether, this data opens the possibility that the consequence of phosphorylation-induced recruitment of c-CBL is the inactivation of the phosphorylated ROR1, similar to a described c-CBL function in other RTKs targeted by c-CBL^31^.

### Lyn KO cells display increased surface levels of ROR1

Our findings reported in Figs. 1 – 4 showed that Lyn can efficiently phosphorylate ROR1 that can be subsequently recognized by c-CBL. ROR1 and Lyn are important regulators of signaling pathways driving chronic lymphocytic leukemia (CLL)^1,8,21,35^ and several lymphomas, namely mantle cell lymphoma (MCL)^15^. Both, Lyn and ROR1, have been evaluated as therapeutic targets in these malignancies and the understanding of their mutual cross talk is thus of direct therapeutic relevance. In order to find out if Lyn controls ROR1 biology in CLL/MCL we decided to generate Lyn-deficient Maver-1 cells. Maver-1^36^ are of MCL origin, express high levels of ROR1 and Lyn, and respond well to BCR activation^37^. We produced Maver-1 Lyn knockout (KO) cells using the Crispr-Cas9 system. We validated four independent clones of Maver-1 Lyn KO cells by western blotting (**Fig. 5a)** and sequencing (Supplementary Table 1). Functionally, Lyn KO Maver-1 cells are deficient in the activation of BCR signaling triggered by IgM. As shown in **Fig. 5b**, activation of several BCR components downstream of Lyn, namely Syk, PLCγ, PI3K, as well as phosphorylation of Lyn substrate HS1 was dramatically reduced in Lyn KO cells. These observations are in line with what is already known about the role of Lyn in the BCR signaling cascade^18^.

**Figure 5:**
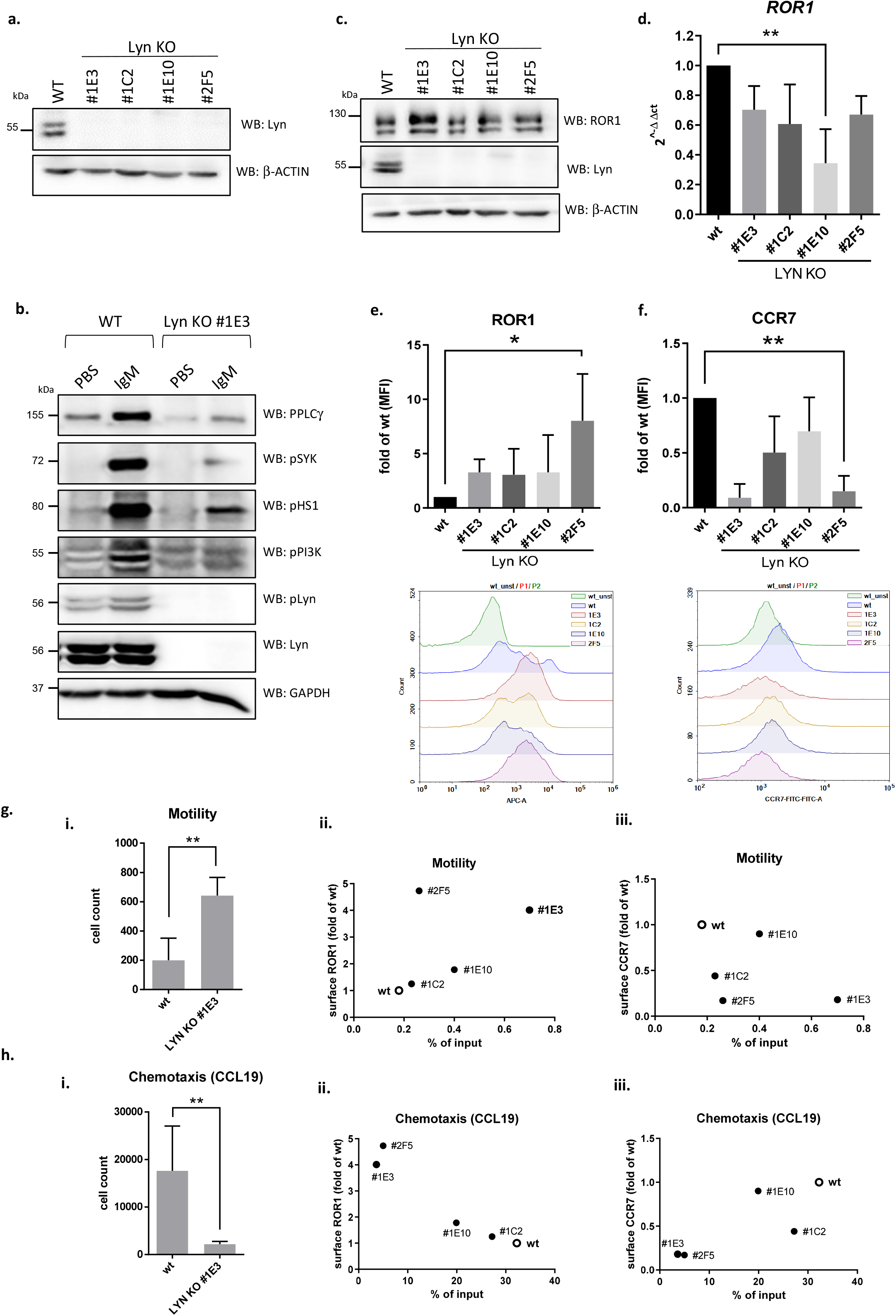
Lyn KO cells display increased surface levels of ROR1. Lyn gene in Maver-1 cells was targeted by Crispr/Cas9. Parental WT and four clones of Lyn KO cells coded #1E3, #1C2, #1E10, and #2F5 were analyzed functionally. **a**) Western blot analysis of expression of Lyn in wt and KO cells. **b**) Activation of BCR signaling after incubation with IgM antibody in WT and Lyn KO (#1E3) cells was assessed by Western blotting of pPLCγ, pSYK, pLyn, pHS1, pPI3K and Lyn. GAPDH was used as a loading control. **c**) Expression of ROR1 protein in WT and Lyn KO clones was analyzed by Western blotting. ß-actin was used as a loading control. Representative blots from 3 independent experiments. **d**) Expression of *ROR1* mRNA in WT and Lyn KO cells. Graphs show mean ± SD of 3 independent experiments. Differences were analyzed by Kruskal-Wallis test (Dunn’s multiple comparison test). **e, f**) Basal surface expression of ROR1 (**e**) and CCR7 (**f**) in Maver-1 WT and individual clones of Lyn KO cells was analyzed by flow cytometry. Mean ± SD from 3 independent experiments and Kruskal-Wallis test (Dunn’s multiple comparison test). Representative histograms for all analyzed clones are shown. **g, h**) Basal motility (**g**) or chemotaxis towards CCL19 (**h**) of Maver wt and Lyn KO cells was analyzed in a transwell assay. Number of cells that migrated to lower chamber of transwell plate after 3 hours of incubation is indicated. Panel i shows mean ± SD from 6 independent experiments for WT and KO clone #1E3, Mann-Whitney nonparametric test. Other panels show variability among 4 different clones; migratory properties are plotted with the surface expression of ROR1 (panel ii) and CCR7 (panel iii) in Maver-1 WT and Lyn KO cells.

After this initial characterization of the BCR signaling deficiency in Lyn KO cells we focused on their ROR1 status. As shown in **Fig. 5c,** global ROR1 protein levels determined by Western blotting were comparable in WT and Lyn KO cells, despite the fact that ROR1 mRNA expression, determined by qPCR, was slightly lower in the Lyn KO cells (**Fig. 5d**). However, when we assessed cell surface ROR1 by flow cytometry we could observe a consistent increase in surface ROR1 in all four clones (**Fig. 5e**). This suggested that endogenous Lyn controls ROR1 trafficking or endocytosis to reduce ROR1 availability on the surface.

ROR1 and its ligand Wnt-5a were shown to control CLL cell migration and chemotaxis^38,39^. More specifically, Wnt-5a-ROR1 axis increases basal migration and reduces chemotaxis^38^. Surprisingly, Lyn KO Maver-1 cells showed not only increased cell surface ROR1 (**Fig. 5e**) but also much lower levels of CCR7 (**Fig. 5f**), a receptor for chemokine CCL19 and essential component of CCL19-induced chemotaxis^40^. Despite some variability in the individual Lyn KO clones, the observed trends showing higher ROR1 and lower CCR7 were always the same (**Fig. 5 e/f**, histograms in the bottom part of the panel).

To address whether the changes in ROR1 and CCR7 levels translate into the changes in the migratory behavior of WT and Lyn KO Maver-1 cells we used transwell assays without (basal motility) or with CCL19 in the bottom chamber (chemotaxis). Lyn KO cells had significantly higher basal motility (**Fig. 5g, i**). This feature was preserved across individual Lyn KO clones and reflected variability of the cell surface ROR1 (**Fig. 5g, ii**) and the cell surface CCR7 (**Fig. 5g, iii**). On the other hand, Lyn KO cells showed reduced chemotaxis to CCL19 (**Fig. 5h, i**) that correlated negatively with cell surface ROR1 (**Fig. 5h, ii**) and positively with the cell surface CCR7 (**Fig. 5h, iii**). Altogether, the analysis of the migratory properties of Lyn KO Maver-1 cells suggested that Lyn, via control of cell surface levels of ROR1 and CCR7, controls the migratory modes of lymphoid cells. Specifically, it suggests that Lyn controls a balance between two migration modes - Wnt-5a/ROR1-mediated basal migration (attenuated by Lyn) and CCL19/CCR7-mediated chemotaxis (promoted by Lyn).

### Cell surface ROR1 is upregulated during CLL migration

Functional experiments with Lyn KO cells indicate that cell surface ROR1 can be under a dynamic control during cell migration. In order to test whether this applies also to primary CLL samples we performed a set of experiments in the patient cohort specified in **Fig. 6a**. Cells from these patients were analyzed by Western blotting by probing for phospho-Y396-Lyn, which is auto-phosphorylated and as such represents a good hallmark for Lyn activation (**Fig. 6b**). p-Lyn levels positively correlated with the phosphorylation of HS1 on tyrosine 397, which is a well described target of Lyn^41^ (**Fig. 6c**). On the other hand, ROR1 levels assessed by Western blotting did not correlate with active Lyn (**Fig. 6d**).

**Figure 6:**
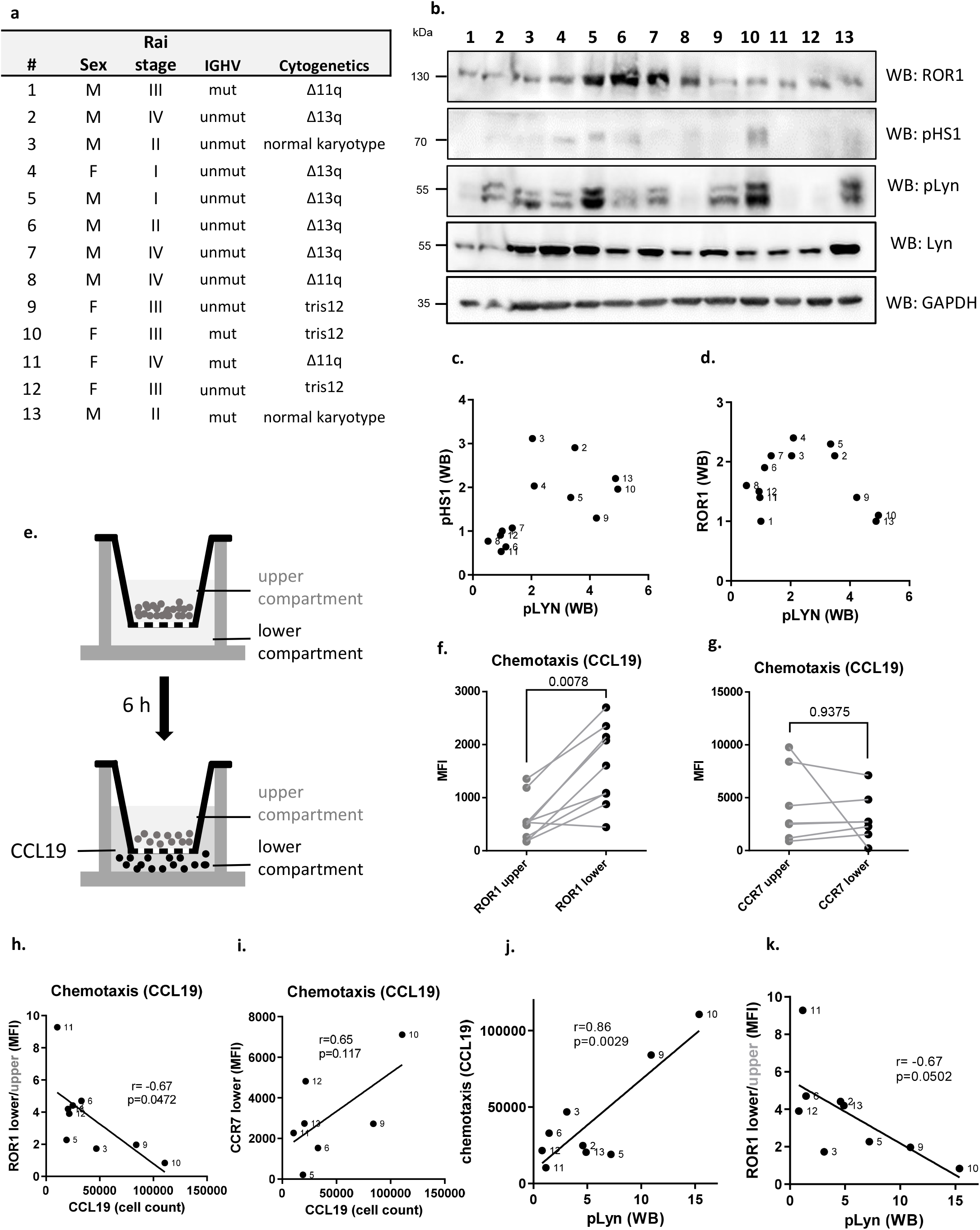
Correlation of Lyn activity with the chemotactic properties of primary CLL cells. **a**) Table with basic clinical characteristics of the patient cohort used for functional analysis of the primary CLL cells. Rai stage at the time of sampling, IGHV status and cytogenetic analysis are indicated (n=13). **b**) Protein levels of ROR1, pLyn, Lyn and pHS1 expression in the panel of CLL primary cells was analyzed by Western blotting. **c, d**) Correlation of pLyn expression with pHS1 (**c**) and ROR1 (**d**). **e**) Scheme of the transwell assay indicating the upper and lower compartment that was used for the separate analysis of the surface markers. **f,g**) Surface expression of ROR1 (f) and CCR7 (g) in the upper (•) and lower (•) compartments of transwell chamber in the panel of primary CLL cells was analyzed by flow-cytometry. Wilcoxon matched pairs signed test (n=9). **h, i**) Correlation of the change in the surface expression of ROR1 (h) and CCR7 (i) during migration represented as the ratio of receptor levels in the lower:upper compartment with the chemotactic properties of cells (expressed as the number of cells in the lower chamber) in the panel of primary CLL cells. **j**) Correlation of the chemotaxis towards CCL19 with pLyn levels (WB). **k**) Correlation of ROR1 surface levels dynamics (expressed as the ratio of ROR1 surface levels in the lower:upper well of transwell) with pLyn levels (WB). **h-k**) Pearson correlation coefficient.

Chemotactic properties of CLL cells were analyzed in transwell assays as the migratory response to the chemokine CCL19. In parallel, we analyzed the surface levels of ROR1 and CCR7 in the non-migratory (upper chamber) and migratory (lower chamber) CLL cells (for schematics see **Fig. 6e**). In some cases, because of low number of migrating cells we were unable to assess in parallel all the functional parameters (migration, CCR7 and ROR1 levels in lower and upper chamber, pLyn levels; see Supplementary Table 2 for details) and as such the number of samples in individual analyses presented below slightly varies. Surprisingly, ROR1 surface expression was clearly increased in the cells that passed through the transwell membrane (**Fig. 6f**). No such increase was observed for CCR7, which is a CCL19 receptor (**Fig. 6g**). Interestingly, the CLL samples that upregulated ROR1 most efficiently, were the least chemotactic (**Fig. 6h**). CCR7 showed an opposite behavior to ROR1 and CCR7 levels correlated positively with the chemotaxis (**Fig. 6i**). This data shows that even in primary CLL cells, similar to Maver-1 cells, ROR1 surface levels are dynamically regulated during cell migration and that ROR1 and CCR7 show opposite behavior, in this respect.

It remained to be analyzed whether the balance between ROR1-driven and CCR7-controlled migration can be correlated with the activity of Lyn in primary CLL cells. pLyn-high CLL samples (based on Fig. 6b), were in general more chemotactic (**Fig. 6j**), which is in line with the positive role of Lyn in chemotaxis identified in Maver-1 cells. On the other hand, pLyn levels negatively correlated with the capacity of CLL cells to upregulate ROR1 during migration (**Fig. 6k**). Altogether, this suggests that the function of Lyn that switches between ROR1- and CCR7-mediated migratory modes, uncovered in Maver-1 cells, is conserved also in primary CLL cells.

## Discussion

A significant amount of research and preclinical development is being conducted on developing monoclonal antibodies to target surface ROR1 in CLL and other malignancies^11,42,43^. Also, a considerable body of work has helped us to understand signaling on the extracellular side of ROR1 via Wnt5a in CLL cells^39,44^. However, significant gaps of knowledge remain in our understanding of the importance of the intracellular domains of ROR1, especially the TKD, as well as of the regulation of ROR1 levels on the cell surface. Our study is the first to show that the Src family kinase Lyn, an important component of BCR signaling, phosphorylates ROR1 intracellularly and controls its surface levels.

The phosphorylation of ROR1 by Lyn identifies a novel crosstalk between ROR1 and BCR signaling. This crosstalk can be of particular importance in CLL and MCL where both, ROR1 and BCR pathways, represent therapeutic targets. It has been shown earlier that in these malignancies the non-canonical Wnt pathway and BCR signaling can be targeted in a combinatorial manner. Namely, it has been observed *in vivo* in the mouse models of CLL where BTK inhibitor ibrutinib and anti-ROR1 antibody^45^ or casein kinase 1 (CK1) inhibitors^46^ showed synergistic effects. Similar behavior has been observed by Karvonen and colleagues in the in vitro model of MCL^15^. Our data suggest that active BCR (correlating with high Lyn activity) negatively controls the surface ROR1. This is an interesting observation in the context of the recent report showing that in MCL ROR1/CD19 membrane complex can functionally compensate for BCR/BTK activity and activate pro-survival and pro-proliferative PI3K-Akt and MEK-Erk cascades^17^. Lyn action towards ROR1 can explain how BCR inhibited cells with low Lyn activation “switch” to the survival mode dependent on ROR1. In addition, study by Zhang et al. opens the possibility that ROR1 and BCR-centered complexes in MCL and CLL share even more components than Lyn described in this study.

Our analysis of Lyn KO Maver-1 cells has identified different migratory properties in comparison to WT cells. Lyn KO cells had higher motility in the absence of external stimuli, a feature correlating with the increased cell surface ROR1, but failed to respond to physiological chemotactic stimulus CCL19, a feature correlating with the decrease in CCR7, a CCL19 receptor. Of interest, similar behavior – i.e. deregulated motility and decreased chemotaxis - was described for aggressive CLL characterized by high Wnt5a expression^38^. This opens the possibility that basal motility, promoted by Wnt5a/ROR1, and chemotaxis represent distinct migratory modes that are coordinated by Lyn activity. In vivo phenotype of Lyn KO leukemic lymphocytes in the TCL1 mouse model, which were more efficient in the spleen infiltration than wt CLL cells^35^ supports this view.

We demonstrate that at least one consequence of Lyn-induced ROR1 phosphorylation is the recruitment of c-Cbl. c-Cbl, a member of a family of RING finger E3 ligases, has been shown to be upregulated in CLL^47^. Out of three 3 different family members - Cbl (a.k.a c-Cbl or RNF55), Cbl-b (RNF56) and Cbl-c (RNF57), Cbl and Cbl-b are known to be highly expressed in B and T lymphocytes. It is tempting to speculate that Lyn activity towards ROR1-induced migration is not limited to the regulation of the interaction with c-CBL but includes also action towards other cytoskeletal modulators of migration such as HS1 and cortactin. Both these proteins serve in CLL cells as substrates of Lyn^41,48^ and at the same time were found to dynamically interact with ROR1 and control ROR1-induced migration^44,49^. HS1-deficient leukemic cells in the mouse model of CLL are more aggressive compared to Lyn wt mainly due to preferential homing to bone marrow^50^ - the molecular mechanism is not known but loss of Lyn capacity to control Wnt5a-driven migration is one possible explanation.

In addition to Lyn, ROR1 has been shown to get phosphorylated also by other TK’s, namely two receptor TK (RTK)s - Met and Src^20^, and MuSK^51^. In the study by Gentile et al^20^, it was shown that ROR1 is first phosphorylated by Met kinase in its PRD and this helps recruit Src which then leads to the phosphorylation of ROR1 in the kinase domain. It remains to be tested whether some membrane associated TKs, such as Axl^52^ or ZAP70^53^ can synergize with Lyn in the regulation of ROR1.

In summary, our study is the first to show the interaction between ROR1, important BCR kinase Lyn and c-Cbl. Our work also provides a molecular mechanism of the crosstalk for two signaling pathways essential for CLL: BCR signaling and the non-canonical Wnt pathway. This crosstalk mechanism provides a basis for the rational combinational therapies targeting BCR and non-canonical Wnt in CLL and MCL.

## Acknowledgement

VB, PK and ZZ gratefully acknowledge the support of the Czech Science Foundation (the projects GA17-09525S, 17-16680S, GA19-20123S). ZD was supported by the European Union Grant FP7 Marie Curie ITN 608180 ‘Wntsapp’. CIISB (LM2018127) and NCMG (LM2015091) research infrastructures funded by MEYS CR are acknowledged for the financial support of the measurements at the Proteomics and Genomics Core Facilities. Further supported by Ministry of Education, Youth and Sports of the Czech Republic (National Program of Sustainability II projects LQ1605 and LQ1601), by the Ministry of Health of the Czech Republic (FNBr 65269705) and by European Structural and Investment Funds, Operational Programme Research, Development and Education – “Preclinical Progression of New Organic Compounds with Targeted Biological Activity” (PreclinProgress; CZ.02.1.01/0.0/0.0/16_025/0007381).

