## Supplementary methods for "Lyn controls chemotaxis and motility of CLL cells via phosphorylation of ROR1"

**Methods**

**Transfections and Treatments**

Transient transfections of HEK-293T cells were carried out using polyethyleneimine (PEI 1 mg/ ml); for transfections in a 10cm plate, a total of 6 µg of DNA and for the 24 well plate a total of 0.2 µg of DNA per well was used. The DNA:PEI ratios were kept at 1:3 (w/v) in all cases. The transfection mixtures were added to cell cultures and incubated for up to 4h after which all medium was aspirated off the cells and they were supplied with fresh DMEM. The cells were harvested for immunoprecipitation or immunocytochemistry 24h post-transfection. To inhibit the kinase activity of Lyn, Dasatinib (Santa Cruz Biotech #218081) was used as indicated. The following plasmids were used for transfection:

All the Lyn expression plasmids were a kind gift from Dr Naoto Yamaguchi (Kasahara *et al.*, 2004). For the expression of hROR1 and tagging, human ROR1 was cloned to pcDNA3.1/V5-His-TOPO. Alternatively, the following plasmids provided by Dr Paolo Comoglio and described previously^20^ have been used: hROR1 cloned into the pRRL2 backbone with no tags; ROR1 intracellular deletion mutants. The mutants hROR1 Y641F, hROR1 Y645F, hROR1 Y646F and hROR1 Y645/646F were generated using the QuickChange II XL site-directed mutagenesis kit (Agilent Technologies #204180a) based on the hROR1 with no tags on the pRRL2 backbone construct. The mutations were verified by sequencing. The primers to create ROR1 point mutants were as follows:

>Y641F Ffwd: GGGCTTTCCAGAGAAATT**TTC**TCCGCTGATTACTACAGG

>Y641F Rev: CCTGTAGTAATCAGCGGAGAAAATTTCTCTGGAAAGCCC

> Y645F Fwd: GAAATTTACTCCGCTGAT**TTC**TACAGGGTCCAGAGTAAGTCC

>Y645F Rev: GGACTTACTCTGGACCCTGTAGAAATCAGCGGAGTAAATTTC

> Y646F Fwd: GAAATTTACTCCGCTGATTAC**TTC**AGGGTCCAGAGTAAGTCC

>Y646F Rev: GGACTTACTCTGGACCCTGAAGTAATCAGCGGAGTAAATTTC

>Y645/646F Fwd	GAAATTTACTCCGCTGAT**TTCTTC**AGGGTCCAGAGTAAGTCC

>Y645/646F Rev: GGACTTACTCTGGACCCTGAAGAAATCAGCGGAGTAAATTTC

**Immunoprecipitation & Western Blot**

For the immunoprecipitation experiments, transfected cells were first washed in cold PBS and then lysed in cold 0.5% NP-40 Lysis buffer (150mM NaCl, 50mM Tris pH 7.6, 1mM EDTA, 0.5% NP40) pH adjusted to 7.5 at 4°C. Prior to lysis, this buffer was supplemented with 1mM Na_3_VO_4_, 1mM DTT, 1mM NaF, cOmplete^TM^ protease inhibitor cocktail and phosphatase inhibitor cocktail set II (Merck). Post incubation with lysis buffer, the cell lysates were collected by gentle pipetting and spun at >14000g for 20 minutes at 4°C. Subsequently, the cleared lysates were transferred to new tubes and used for protein concentration measurement using the DC^TM^ protein assay kit (BIO-RAD). 500 µg of total protein was taken for the immunoprecipitation and incubated with 1 µg of antibody with gentle rotation over-night, followed by incubation with Protein G Sepharose beads (GE Healthcare, #17–0618-05) or Dynabeads^TM^ (ThermoFisher Scientific #10003D) for 2-4 h. The beads were then washed 4 times in cold lysis buffer and after the last wash resuspended directly in 2x Laemmli buffer. For immunoprecipitation, the antibodies used were: ROR1 (RnD AF2000), Normal goat IgG (RnD #AB-108-C), Lyn (Novus Biologicals #NB-500-519), Normal mouse IgG (Merck #12-371), V5 (Life Technologies # R96025) and pY (Millipore #05-321).

Primary CLL cells and Maver-1 cells were for western blotting lyzed in 1% SDS buffer (1% (w/v) SDS, 10 mM TRIS, 1 mM EDTA, pH 8.0) supplemented with cOmplete^TM^ protease inhibitor cocktail and phosphatase inhibitor cocktail set II (Merck). For the western blotting, samples were briefly boiled at 95°C for 5’ and then loaded on 8% gels and separated by SDS-PAGE. The transfer step was carried out on to Immobilon-P^®^ (Merck) PVDF membranes at 106 V for 75 minutes. Membranes were blocked for an hour at room temperature in 3% non-fat milk or BSA prepared in TBS/T buffer. The antibodies used for WB were diluted in the TBS/T buffer containing 5% BSA and 0.02% NaN_3_.

The primary antibodies used were as follows: ROR1 (CST #4102), anti-pY (Millipore #05-321), anti- V5 (Life Technologies #R96025), anti-CBL (CST #8447), Lyn monoclonal (CST #C13F9), Lyn polyclonal (CST # 2732), Lyn (Novus Biologicals NB500-519), phospho-Y396 Lyn (Abcam ab226778), pPLC g2 (CST #3874), pSYK (CST #2710), pBTK (CST #5082), phosho-Y397 HS1 (CST #4507), pPI3K (CST #4228S), GAPDH (CST #5174S), β-Actin (CST #4970). In addition, hROR1 antibody which was a kind gift from Dr. Henry-Hsin Ho was also used. Post-secondary antibody incubation with Goat Anti-Rb IgG (whole molecule) – Peroxidase antibody (#A0545 Sigma-Aldrich), Clean Blot IP detection reagent HRP (Thermo Scientific #21230) or Goat Anti-Mouse IgG (whole molecule) – Peroxidase antibody (#A4416 Sigma-Aldrich), the membranes were incubated with Immobilon western chemiluminescent HRP substrate (Merck) and developed using the FusionSL imaging system (Vilber-Lourmat).

**Immunofluorescence:**

In a 24 well plate, 0,05*10^^6^ HEK-293T WT cells per well were seeded on coverslips which were pre-treated with 0.1% gelatin or 1:10 diluted poly-L-Lysine and transfected the following day. After 24 hours, the cells were washed with PBS and then fixed in 4% paraformaldehyde for 15 min at R.T. The cells were then washed in PBS (3x) and treated for 1h with PBTA buffer (1x PBS, 5% BSA, 0.25% Triton X-100, 0,02% NaN_3_) to simultaneously block as well as permeabilise the cells. Next, the cells were incubated over night at 4°C with the primary antibodies diluted (1:500) in PBTA buffer. On the following day, the cells were washed in PBS (3x) and treated with secondary antibodies conjugated to Alexa fluorophores for 2h. From this point on, the 24 well plate was protected from light. Finally, the wells were washed with PBS (3x) and for the 4^th^ wash were treated with DRAQ5 in PBS (1:2000) for 5-10 minutes to stain the nuclei. The DRAQ5 (CST #4084S) solution was pipetted off and the cells were washed one last time in PBS. The coverslips were mounted onto microscopy slides using glycerol-gelatin mounting medium (Sigma Aldrich #GG1). Leica SP8 confocal microscope was used to visualize the cells under the 60x (Oil) objective. The primary antibodies used were as follows: ROR1 (RnD AF2000) and Lyn (Novus Biologicals NB500-519). The secondary antibodies used were as follows: Alexa 488 donkey anti-goat (Invitrogen A-11055), Alexa 568 donkey anti-mouse (Invitrogen A-10037).

**CRISPR/Cas9 gene editing in Maver-1 cells**

***Lentiviral infection***

Lyn KO Maver-1 clone 1E3 was produced using lentiviral infection. lentiGuide-Puro (Addgene plasmid # 52963; http://n2t.net/addgene:52963; RRID:Addgene_52963) and lentiCas9-Blast (Addgene plasmid # 52962; http://n2t.net/addgene:52962; RRID:Addgene_52962) were a gift from Feng Zhang. To produce lentiviruses the transfer plasmid (lentiGuide-Puro) was co-transfected with the viral envelope vector pCMV-VSV-G (Addgene #8454) and packaging vector pCMV.dR8.91 (kind gift of Dr. Didier Trono).

To target exon 4 of Lyn, the following pair of annealed oligos were cloned into the single gRNA scaffold of lentiGuide-Puro: Oligo1-CACCGTGAAAGACAAGTCGTCCGGG, Oligo2: AAACCCCGGACGACTTGTCTTTCAC

*Viral particle production:* HEK-293 cells (ATCC® CRL-1573) were seeded into 6-well plates (0.4 x 10^6^ cells per well). The plasmids were transfected using PEI and Opti-MEM. For the transfection, we used a DNA:PEI ratio of 1:2 (w/v). The mix of plasmids for viral production also included the pCMV-GFP (Addgene #11153) to check the efficiency of transfection. The transfection was allowed to continue for up to 3 h before replacing the medium with fresh 1 ml DMEM in each well. Viral particles were harvested on Day 2 and 3 post-transfection; on each day there was an interval of 8h between the first and second harvest. Each time the medium was carefully aspirated off the cells into 50 ml falcon tubes and fresh 1ml complete medium was supplied to the cells. After the final harvest, the medium with the viral particles was aliquoted (1 ml) into eppendorf tubes and stored at -80°C.

*Transduction and selection:* Maver-1 cells (1x10^6^) were used for the transduction in 2 steps. In the first step, the cells were transduced to express Cas9. Briefly, 1 µl of Polybrene (Merck Cat No: TR-1003-G) was added to 1 ml of the Cas9 expressing virus to a final concentration of 10 ug/ml. This mix was then added very gently to a pellet of the Maver-1 cells and mixed well to resuspend the cells. The cells were then centrifuged for 30 min at 800 g at 37°C. Post centrifugation, the pellet was once again resuspended into the same medium and the cells were seeded into 24 well plates. After 24 h, these cells were centrifuged and the medium containing the virus and polybrene was aspirated off. The cells were washed once in PBS and then finally resuspended in fresh RPMI, the optimum medium for Maver-1 cells. Blasticidin (Invivogen #ant-bl-05) was added (10 µg/ml) and the transduced cells were allowed to undergo selection for 4 days, after which they were spun down and then resuspended in new medium with fresh Blasticidin for an additional 4 days. As controls, we used non-transduced Maver-1 cells supplied with medium containing only polybrene. Cas9 expressing cells (as confirmed by WB – CST #14697) were then subjected to a second round of transduction with the virus carrying the gRNA, the same way as described above. After 24 h of transduction, these cells were washed in PBS and resuspended in fresh medium supplied with Puromycin (Invivogen #ant-pr-1) at a final concentration of 10 µg/ml. The cells were allowed to undergo selection for 4 days. The surviving polyclonal population was passaged for 2 additional rounds and further subjected to the limiting dilution method to obtain a monoclonal population of cells.

***Production of Maver-1 KO cells by electroporation***

All other Maver-1 Lyn KO clones except 1E3 were produced by electroporation using the Neon Transfection System (Invitrogen). We used the same gRNA as in the lentiviral infection, but this time cloned into an all-in-one expression vector pSpCas9(BB)-2A-GFP (PX458) (Addgene #48138). The cells (2.5 million/reaction) first underwent 2 cycles of PBS wash and centrifugation for 5 minutes at 200 g and then were resuspended again in 100 µl of PBS. The cell suspension was then mixed with 10 µl of 2µg/µl plasmid solution in MQ water. The mix was then electroporated using Neon and 100µl Neon tips, with the pulse setup of 1600/10/3 (pulse intensity/pulse width/pulse number). Electroporated cells were resuspended in full RPMI medium and incubated overnight. To get the monoclonal populations, GFP-positive cells were sorted as single cells to 96-well culture plates with U bottom containing RPMI medium with 20 % FBS.

**Mass spectrometric Analysis**

***Phospho analysis***

Immunoprecipitated samples were loaded onto SDS-PAGE gels (Coomasie brilliant blue staining). Protein in gel pieces were reduced, alkylated, digested by trypsin and subsequently cleaved by chymotrypsin. Digested peptides were extracted from gels. 1/10 of the peptide mixture was analyzed directly and the rest of the sample was enriched by Pierce Magnetic Titanium Dioxide Phosphopeptide Enrichment Kit (Thermo Scientific, Waltham, Massachusetts, USA) according to manufacturer protocol and eluted into LC-MS vial with already added PEG (final concentration 0.001%). Eluates were concentrated under vacuum and then dissolved in water and 0.6 μl of 5% FA to get 12 μl of peptide solution before LC-MS/MS analysis.

LC-MS/MS analyses of peptide mixture were done using Ultimate RSLCnano system connected to Orbitrap Elite hybrid spectrometer (Thermo Fisher Scientific) with ABIRD (Active Background Ion Reduction Device; ESI Source Solutions) and Digital PicoView 550 (New Objective) ion source (tip rinsing by 50% acetonitrile with 0.1% formic acid) installed. Prior to LC separation, tryptic digests were online concentrated and desalted using trapping column (100 μm × 30 mm) filled with 3.5-μm X-Bridge BEH 130 C18 sorbent (Waters). After washing of trapping column with 0.1% FA, the peptides were eluted (flow 300 nl/min) from the trapping column onto Acclaim Pepmap100 C18 column (3 µm particles, 75 μm × 500 mm; Thermo Fisher Scientific) by 65 min long gradient. Mobile phase A (0.1% FA in water) and mobile phase B (0.1% FA in 80% acetonitrile) were used in both cases. The gradient elution (300 nl/min) started at 1% of mobile phase B and increased from 1% to 56% during the first 50 min (30% in the 35th and 56% in 50th min), then increased linearly to 80% of mobile phase B in the next 5 min and remained at this state for the next 10 min.

MS data were acquired in a data-dependent strategy selecting up to top 10 precursors based on precursor abundance in the survey scan (350-2000 m/z). The resolution of the survey scan was 60 000 (400 m/z) with a target value of 1×10^6^ ions, one microscan and maximum injection time of 200 ms. High resolution (resolution 15 000 at 400 m/z) HCD MS/MS spectra were acquired with a target value of 50 000. Normalized collision energy was 32 % for HCD spectra. The maximum injection time for MS/MS was 500 ms. Dynamic exclusion was enabled for 45 s after one MS/MS spectra acquisition and early expiration was disabled. The isolation window for MS/MS fragmentation was set to 2 m/z. The analysis of the mass spectrometric RAW data files was carried out using the Proteome Discoverer software (Thermo Fisher Scientific; version 1.4) with in-house Mascot (Matrixscience, London, UK; version 2.4.1) search engine utilization. The phosphoRS feature was used for phosphorylation localization. Peptides with false discovery rate (FDR; q-value) < 1%, rank 1 and with at least 6 amino acids were considered. Quantitative information was assessed and manually validated in Skyline software (Skyline daily 3.1.1.8884).

### *Analysis of IP samples*

Immunoprecipitates were digested by trypsin (Promega) on-beads for 2 hours (at 37 ºC), then supernatant was transferred to a new vial and incubated with trypsin overnight (at 37 ºC). The resulting peptides were extracted into LC-MS vials by 2.5% formic acid (FA) in 50% acetonitrile (ACN) and 100% ACN with addition of polyethylene glycol (0.001%) and the sample volume was reduced to 15 μl in a SpeedVac concentrator (Thermo Fisher Scientific). LC-MS/MS analyses of peptide mixture were done using system described in previous section. Prior to separation, tryptic digests were concentrated and desalted on trap column (X-Bridge BEH 130 C18 trap column (3.5 mm particles, 100 mm x 30 mm; Waters) with 0.1% FA, then the peptides were separated on an analytical column (Acclaim Pepmap100 C18, Thermo Fisher Scientific) by 120 min nonlinear gradient program (mobile phase A: 0.1% FA in water; mobile phase B: 0.1% FA in 80% can; 300 nl/min). The gradient elution started at 1% of mobile phase B and increased from 1% to 56% during the first 105 min (30% in the 75th and 56% in 105th min), then increased linearly to 80% of mobile phase B in the next 5 min and remained at this state for the next 10 min. MS data were acquired in same way as in previous case (please see Phosphoanalysis section). The analysis of the mass spectrometric RAW data files was carried out using the MaxQuant software (version 1.6.2.10) using default settings unless otherwise noted. MS/MS ion searches were done against modified cRAP database (based on http://www.thegpm.org/crap) containing protein contaminants like keratin, trypsin etc., and UniProtKB protein database for Homo Sapiens (<ftp://ftp.uniprot.org/pub/databases/uniprot/current_release/knowledgebase/reference_proteomes/Eukaryota/UP000005640_9606.fasta.gz>; downloaded 19.09.2018, version 2018/08, number of protein sequences 21,053). Oxidation of methionine and proline, deamidation (N, Q) and acetylation (protein N-terminus) as optional modification, carbamidomethylation (C) as fixed modification and trypsin/P enzyme with 2 allowed miss cleavages were set. Peptides and proteins with FDR threshold < 1% and proteins having at least one unique or razor peptide were considered only. Match between runs was set among all analysed samples. Protein abundance was assessed using protein intensities calculated by MaxQuant.

Protein intensities reported in proteinGroups.txt (from MaxQuant) were further evaluated using the software container environment (<https://github.com/OmicsWorkflows/KNIME_docker_vnc>; version 3.7.2a). Processing workflow is available upon request. Briefly, it covered following steps: a) removal of decoy hits and contaminant protein groups, b) protein group intensities log_2_ transformation, c) LoessF normalization and d) ratio calculations from normalized protein-group intensities based on the experiment design between controls and ROR samples. Each ratio was compared with defined thresholds for each comparison and the final list of differently expressed proteins was made based on the following criteria: the fold change threshold > 5, protein groups had to have at least 2 peptides and non-zero protein group intensity. UpSet plot was produced using UpSetR R package (version 1.4.0) in R (version 3.6.1).

**Quantitative real-time PCR**

Total RNA was obtained from the cells using the RNeasy Mini Kit (Qiagen), as per the manufacturer’s instructions. 1 µg of RNA was reverse transcribed using dNTP’s mix, oligo dT primers (Fermentas), and SuperScript™ II reverse transcriptase (Invitrogen). The expression levels of *ROR1* and reference gene *GAPDH* were measured using Power SYBR Green master mix (ThermoFisher Scientific).

The primers used were as follows:

h*ROR1:* For: TCTCGGCTGCGGATTAGAAAC, Rev:TCCAGTGGAAGAAACCACCTC

*TBP*: For: GAGAGTTCTGGGATTGTACCG, Rev: ATCCTCATGATTACCGCAGC

*GAPDH:* For: GACAGTCAGCCGCATCTTCT**,** Rev: TTAAAAGCAGCCCTGGTGAC

**Supplementary Table 1 (Related to generation of Lyn KO Maver-1 cell line in Methods)**

| Lyn Gene | **Chromosome 8, (-) strand**  **Screening primers-**  **Fwd**: ATCCTTTGTGCCCCATGAGG  **Rev:** ATTACAAGAAAGGGGCCGGG |
| --- | --- |
| 1E3 KO Seq | 1nt ins (G)  GTGCCCCATGAGGTTTGTTAAAACGCTTCTGCTGATGGATTCTTACAGGTGTTCTCTTGTGTTCATCTTTAGATCCAGAGGAACAAGGAGACATTGTGGTAGCCTTGTACCCCTATGATGGCATCCACCCGGGACGACTTGTCTTTCAAGA |
| 2F5 KO Seq | Allele 1: 1nt del (G)  TTTGTGCCCCATGAGGTTTGTTAAAACGCTTCTGCTGATGGATTCTTACAGGTGTTCTCTTGTGTTCATCTTTAGATCCAGAGGAACAAGGAGACATTGTGGTAGCCTTGTACCCCTATGATGGCATCCACCCGGACGACTTGTCTTTCAAG  Allele 2: 1nt ins (G)  GCCCCATGAGGTTTGTTAAAACGCTTCTGCTGATGGATTCTTACAGGTGTTCTCTTGTGTTCATCTTTAGATCCAGAGGAACAAGGAGACATTGTGGTAGCCTTGTACCCCTATGATGGCATCCACCCGGGACGACTTGTCTTTCAAGAAA |
| 2C10 KO Seq | 1nt ins (G)  GTGCCCCATGAGGTTTGTTAAAACGCTTCTGCTGATGGATTCTTACAGGTGTTCTCTTGTGTTCACCTTTAGATCCAGAGGAACAAGGAGACATTGTGGTAGCCTTGTACCCCTATGATGGCATCCACCCGGGACGACTTGTCTTTCAAGA |
| 1C2 KO Seq | Allele 1: 1nt del (G)  GTGCCCCATGAGGTTTGTTAAAACGCTTCTGCTGATGGATTCTTACAGGTGTTCTCTTGTGTTCATCTTTAGATCCAGAGGAACAAGGAGACATTGTGGTAGCCTTGTACCCCTATGATGGCATCCACCCGGACGACTTGTCTTTC  Allele 2: 1nt ins (G)  CCCCATGAGGTTTGTTAAAACGCTTCTGCTGATGGATTCTTACAGGTGTTCTCTTGTGTTCATCTTTAGATCCAGAGGAACAAGGAGACATTGTGGTAGCCTTGTACCCCTATGATGGCATCCACCCGGGACGACTTGTCTTTCAAGAACG |

**Supplementary Table 2 (Related to Figure 6, main text)**

| **patient #** | **chemotaxis** | **ROR1 lower** | **ROR1 upper** | **CCR7 lower** | **CCR7 upper** | **pLYN (WB)** | **pHS1 (WB)** |
| --- | --- | --- | --- | --- | --- | --- | --- |
| **1** |  |  |  |  |  | x | x |
| **2** | x | x | x |  |  | x | x |
| **3** | x | x | x |  |  | x | x |
| **4** |  |  |  |  |  | x | x |
| **5** | x | x | x | x | x | x | x |
| **6** | x | x | x | x | x | x | x |
| **7** |  |  |  |  |  | x | x |
| **8** |  |  |  |  |  | x | x |
| **9** | x | x | x | x | x | x | x |
| **10** | x | x | x | x | x | x | x |
| **11** | x | x | x | x | x | x | x |
| **12** | x | x | x | x | x | x | x |
| **13** | x | x | x | x | x | x | x |
